# Human Cytomegalovirus Infects and Remodels 3D-biofabricated Skin Equivalents

**DOI:** 10.64898/2026.08.21.746345

**Authors:** Haiwei Zhai, Hunter Novacek, Olivia Warriner, Meghan Biegert, Peter Angeletti, Fanben Meng, Lindsey B. Crawford

## Abstract

Viral infection, including for the prototypical betaherpesvirus human cytomegalovirus (HCMV) is typically studied in two-dimensional monocultures, which provide experimentally tractable systems for measuring viral replication but do not reproduce the biologically accurate multicellular organization or interactions, nor the three-dimensional (3D) architecture of stratified epithelial tissues. Here, we evaluated a biofabricated 3D skin model containing fibroblasts and keratinocytes as a system to study HCMV infection. The model is generated using a fibrin-based matrix and spatially organization deposition of keratinocytes in combination with fibroblasts, followed by cellular differentiation controlled by calcium conditions. Cultures were infected with a GFP-expressing clinical strain of HCMV (TB40/E-GFP) and compared to traditional monolayer cultures or single cell type 3D cultures. HCMV infection was detectable by GFP expression in an MOI-dependent and longitudinal manner. Infectious virus was recovered from both the cellular associated and released into the extracellular space of the model, demonstrating that the model supports productive viral infection. Transit of infectious virus is reduced in both multicellular and single type 3D cultures, suggesting that the matrix composition influences viral kinetics. Treatment with the antiviral Ganciclovir suppressed virus production in both fibroblast monolayer cultures and 3D skin cultures. These findings establish a tractable, longitudinal observable, multicellular 3D system that supports productive HCMV infection and provides a reliable platform for investigating viral replication in a spatially organized tissue context.

## Introduction

Human cytomegalovirus (HCMV) is a ubiquitous human betaherpesvirus that establishes lifelong infection following primary exposure. Productive, lytic, HCMV replication is commonly studied in monolayer cultured fibroblast, which support high-titer virus production and have provided the foundation for numerous mechanistic studies of HCMV replication (1). However, HCMV infects a broad range of cell types in vivo, and infection of epithelial, endothelial, stromal cells, and hematopoietic lineage cells all contribute in cell-type specific manners to viral dissemination and pathogenesis. Monolayer (or single cell type) cultures can also be used to study broader ranges of cell types, but analysis in combination is still limited.

The cellular context in which HCMV infection occurs is therefore an important consideration for understanding the broad view of both viral pathogenesis and host responses. Prior studies have demonstrated HCMV infection in multiple cell types, including adult human skin explants (2), a salivary epithelial model derived from human salivary gland tissue (3), and those derived from induced pluripotent stem cells (iPSCs) to model neurodevelopmental changes in brain organoid systems (4-7).

Despite these advances, most experimental studies of HCMV infection, especially for lytic replication and cellular remodeling, remain dependent on two-dimensional monocultures. Monolayer cultures provide significant experimental advantages including defined cell populations, ease of imaging, molecular manipulation, and quantitative measurement of viral replication. However, they do not reproduce the three-dimensional architecture, extracellular matrix interactions, spatial relationships, multicellular organization of a differentiated tissue, nor cell-to-cell viral spread between cell types; all of which are important for viral entry, spread, cell survival, cellular differentiation, and host responses.

Three-dimensional skin models provide an opportunity to address these limitations while retaining experimental accessibility. Biofabrication approaches can establish spatially defined cellular arrangements and promote organization of cells into multilayered structures. Zhai et. al., previously developed a 3D-bioprinting strategy in which keratinocyte-containing fibrin constructs undergo spatially guided migration and self-organization. These then produce multilayered epidermal structures with basal-to-suprabasal organization and cell-cell junctions (8-10). This system provides a tissue-like environment while permitting use of primary human cells or cell lines, experimental manipulation, longitudinal observation, and replicate experiments using the same cell sources at different times. from the same cell-donor matched replicates.

This study addressed whether a biofabricated multicellular skin model can support experimental HCMV infection. We focused initially on a model containing both fibroblasts and keratinocytes and compared infection with conventional 2D monocultures. We used a GFP-expressing HCMV strain to permit longitudinal visualization of infected cells to determine whether the 3D culture system provides a tractable model in which infection can be established and followed over time. We demonstrate that the biofabricated skin model can be productively infected with HCMV, has similar responses to antiviral treatment, and that infection progression can be measured using traditional techniques.

## Results

### A biofabricated fibroblast-keratinocyte culture supports longitudinal HCMV infection

To determine if HCMV can infect biofabricated skin equivalents, we first generated a fibroblast-keratinocyte multicellular skin model as previously described (8, 9). Briefly, fibroblasts are seeded within a fibrin gel serving as the dermal matrix while keratinocytes are encapsulated and bioprinted into fibrin microdroplets for epithermal assembly. Progressive culture and a switch from calcium-free to high calcium containing media was used to induce keratinocyte differentiation within the system. Skin organ models infected with HCMV (**Figure 1A**) demonstrate relatively normal observed morphological differentiation through a 14-day culture period following infection. To establish reference infection, HCMV infection was examined in parallel in fibroblast and keratinocyte monolayers (**Figure 1B**) and in monocultures with the 3D matrix (**Figure 1B**).

**Figure 1.**
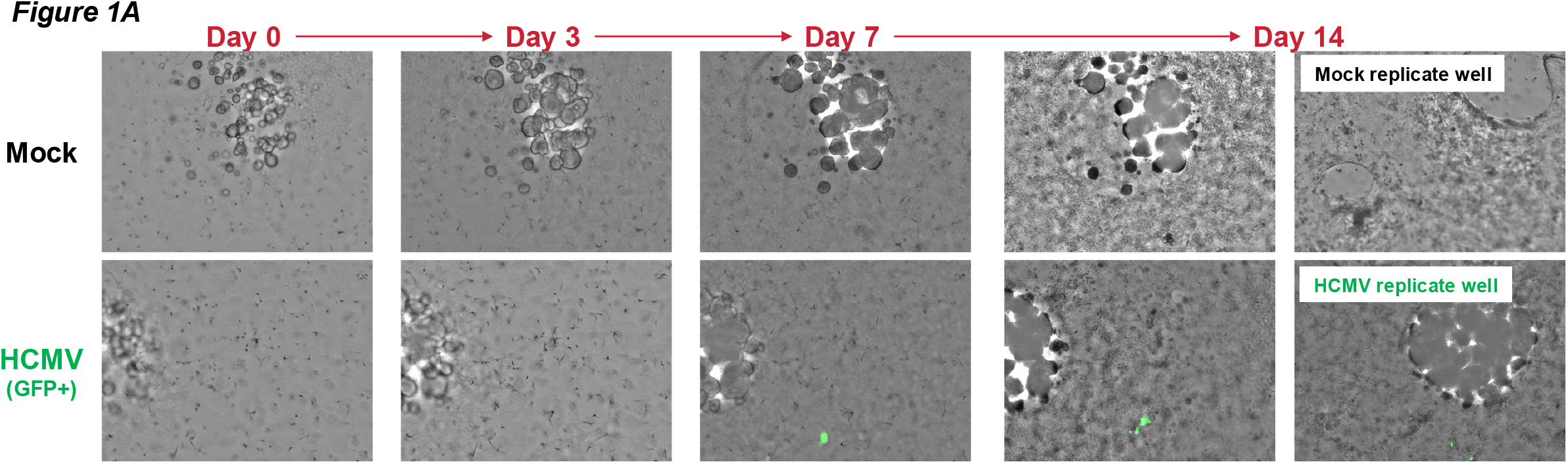
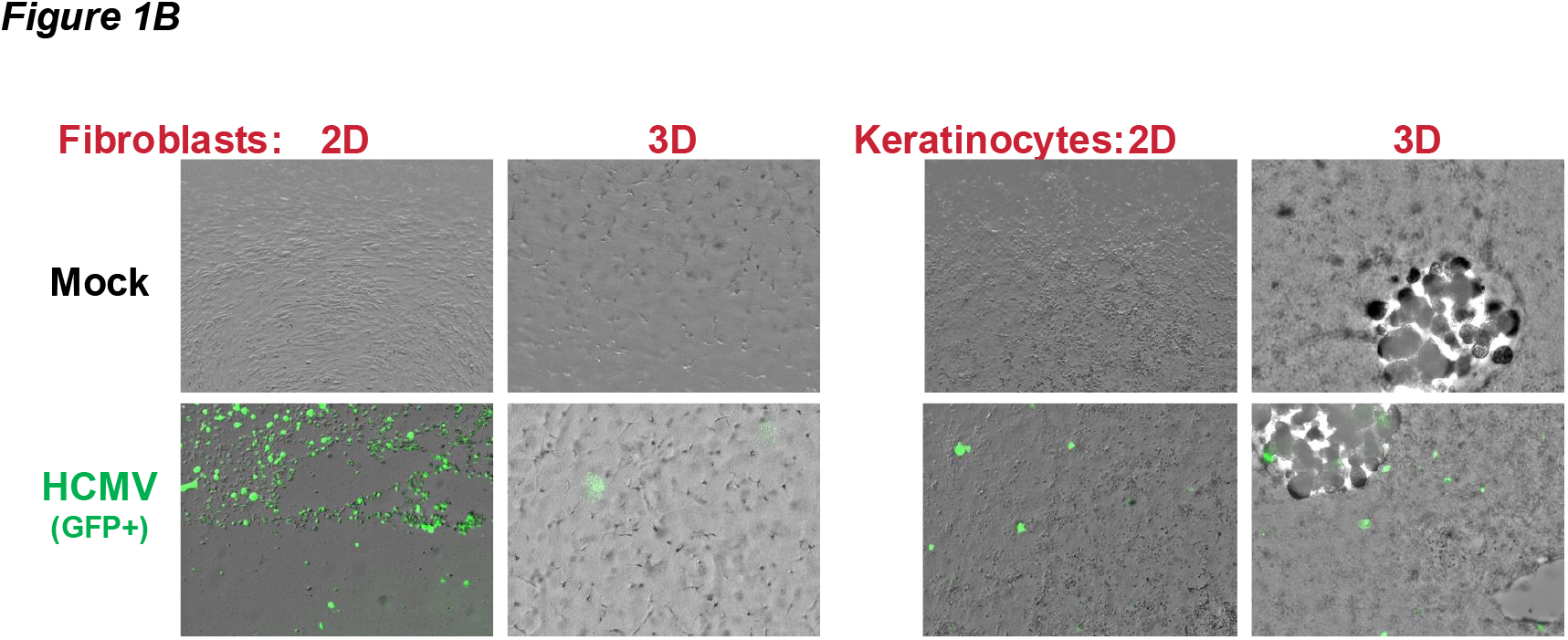
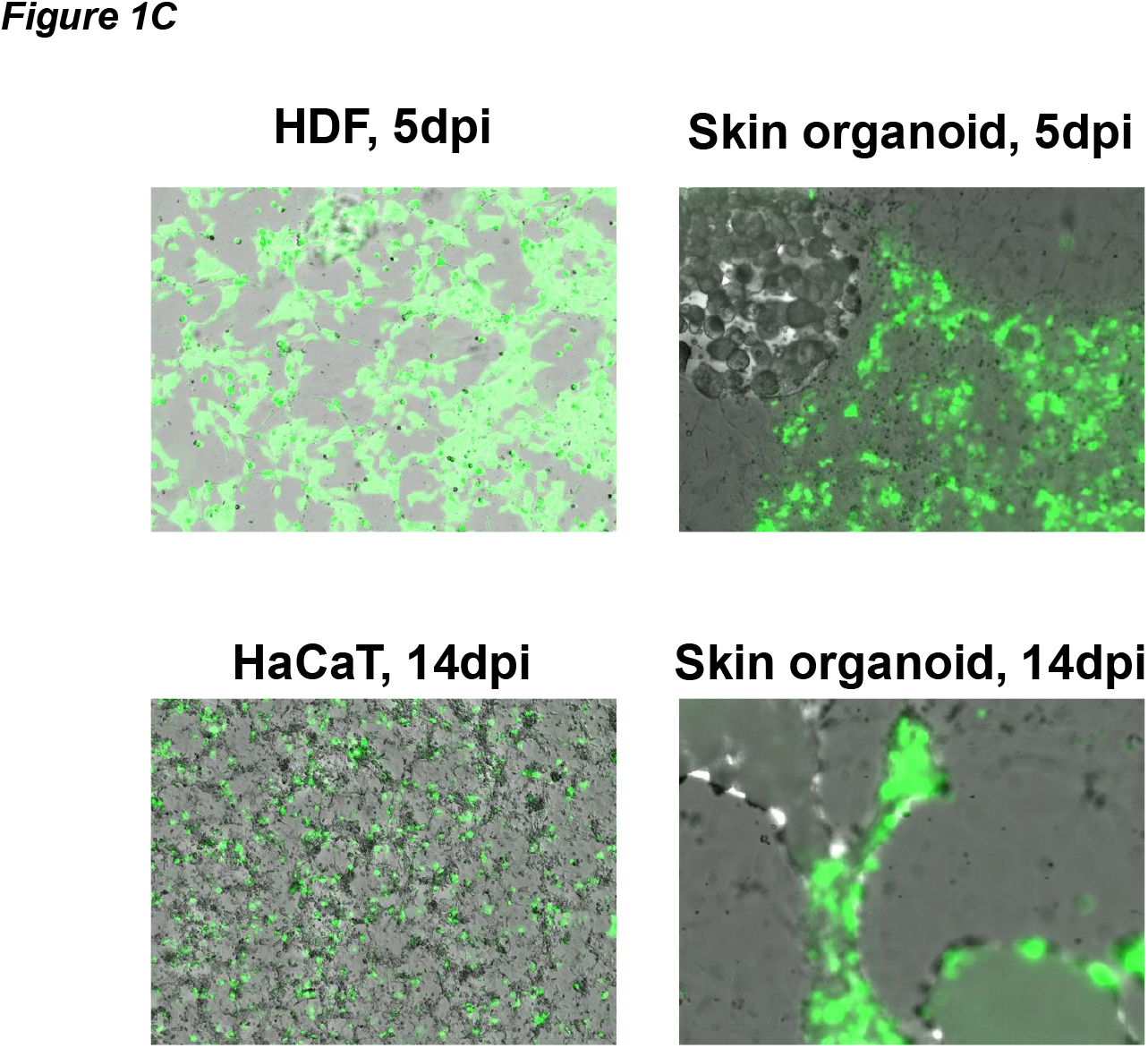
Biofabricated fibroblast-keratinocyte culture supports longitudinal HCMV infection. 3D skin cultures (**A**) were infected with HCMV (day 0) and cultured in parallel with single cell 3D cultures and standard monolayer cultures (**B**). Images were acquired as shown using a SpectraMax Imaging Cytometer (Molecular Devices) allowing remeasurement of the same coordinates on the plate on additional days. HCMV can be visualized by GFP expression as expressed from the virus. Images shown are from one out of four experimental replicates. An additional timecourse is shown in part in **C** and in entirety in Supplemental Figures 1-3.

At low multiplicity of infection (MOI) (**Figure 1A**), skin organ cultures show limited infected cells as determined by viral GFP expression. Fibroblast monolayer cultures (**Figure 1B**), in comparison, demonstrate viral spread and cell loss as expected. The keratinocytes used here also demonstrate permissivity to infection and GFP expression. At higher MOI (**Figure 1C** and **Supplemental Figures 1-3**), an increase in viral infection as measured by observed GFP expression is seen in all cell types.

### 3D skin models support productive HCMV replication

To determine if HCMV infection is productive rather than simply permissive for initial protein expression (e.g., GFP expression), we collected supernatants from infected cultures, including standard monolayer fibroblasts (2D), keratinocyte monolayers (2D), and 3D skin equivalent cultures (**Figure 2A**). Infected fibroblasts demonstrate expected viral replication following infection, while skin organ cultures show reduced infection but do release infectious virus on a delayed timeline.

**Figure 2.**
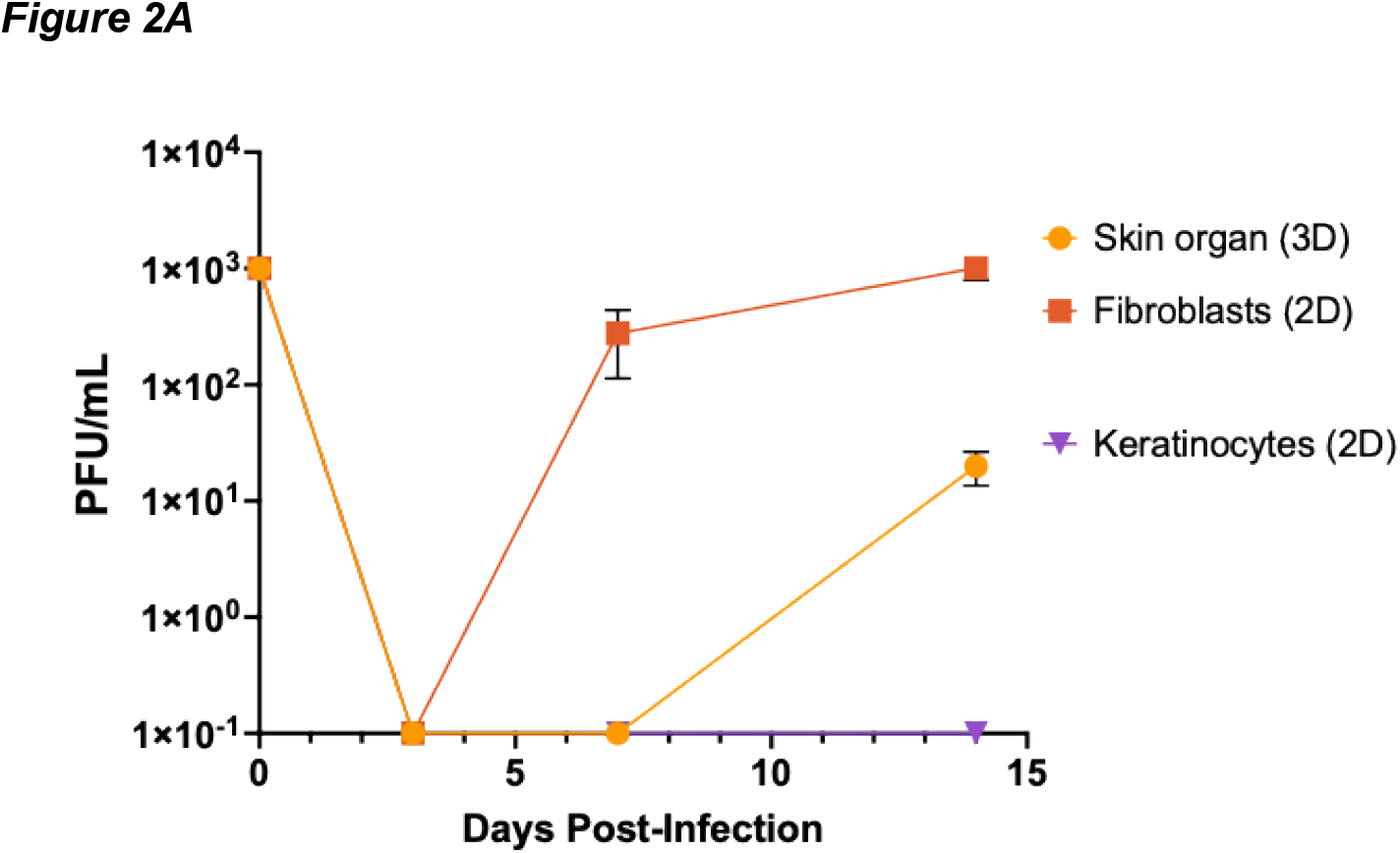
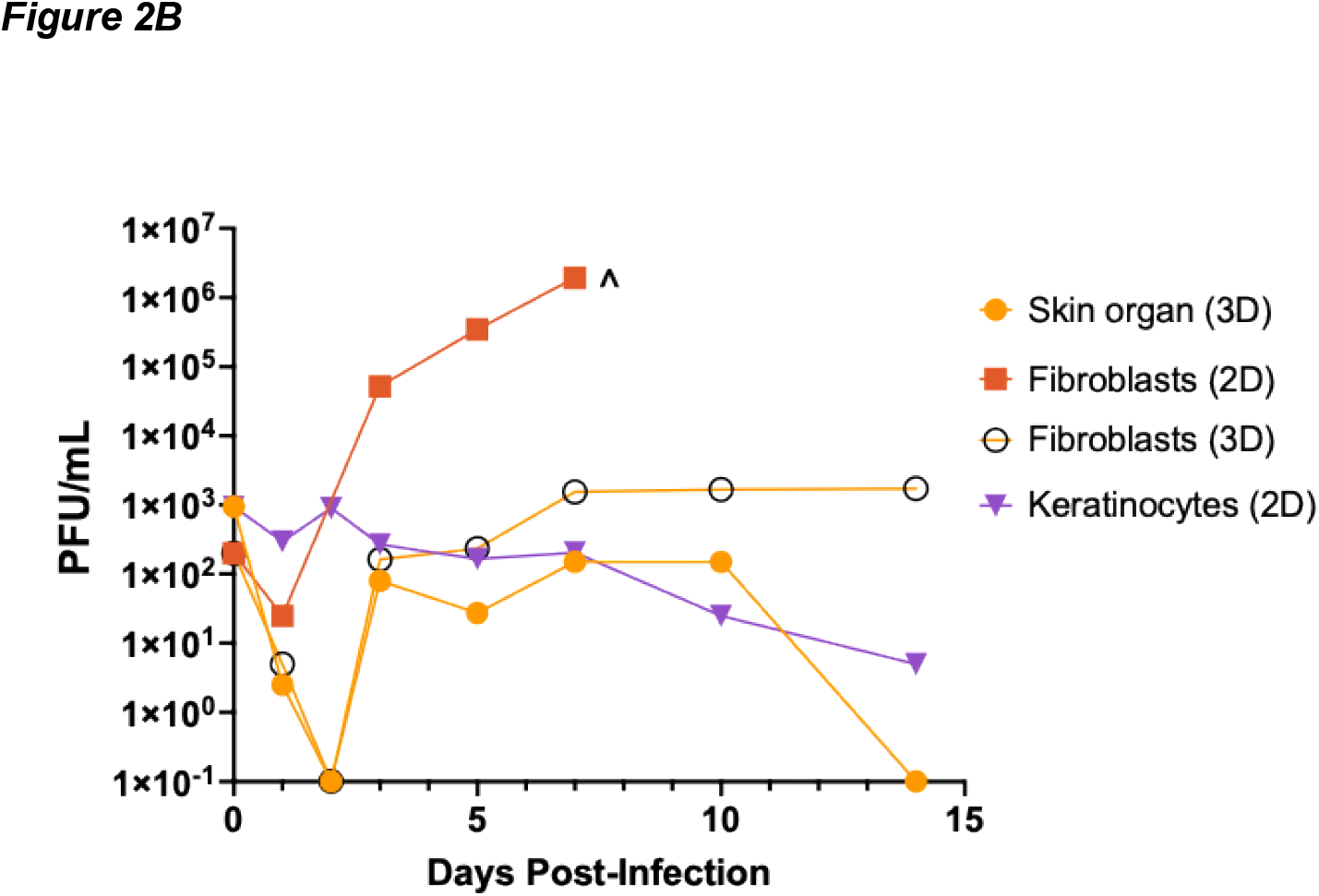
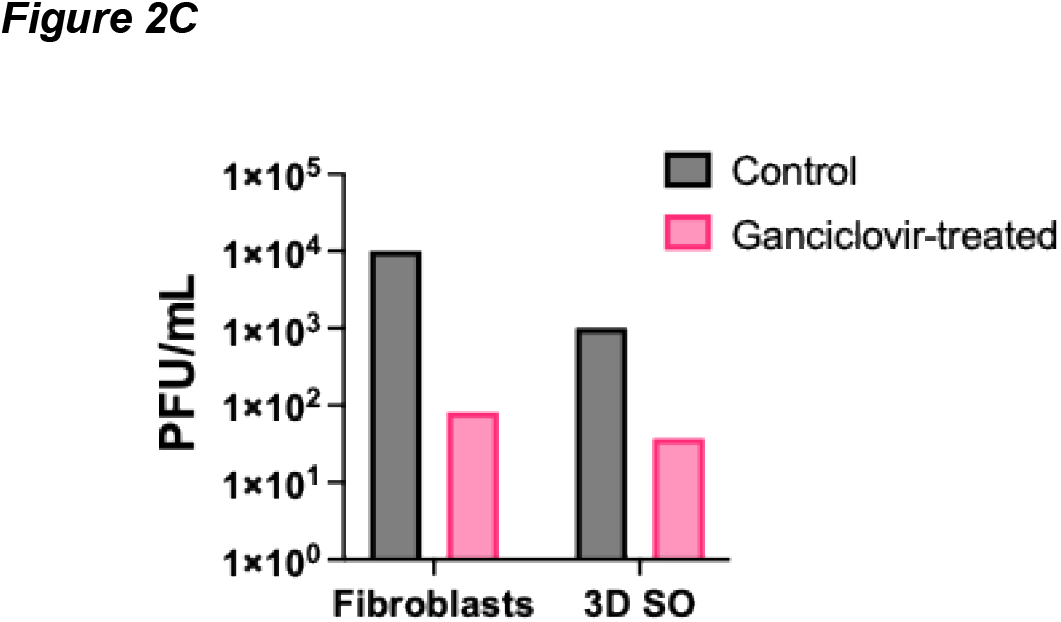
3D skin models support productive HCMV replication with antiviral-induced suppression. Cultures, including: standard fibroblast monolayers (orange, square) and keratinocytes monolayers (purple, inverted triangle), single cell type 3D culture (fibroblasts, yellow, open circle), and 3D skin models (yellow, closed circle) were infected with HCMV and cell-free supernatant harvested at the indicated times. Infectious virus was quantified by standard plaque assay on fibroblasts (**A and B**) with duplicate titer wells. Data shown is the mean +/-standard error of the mean for four independent experiments (**A**). In (**B**), a lower MOI (50% infectious dose) was used in the fibroblast monolayer to control for increased viral replication and the monolayer assay discontinued when the monolayer was destroyed (^). **C)** Replicate infected wells for both the 3D skin organ culture and the standard fibroblast monolayer were treated with Ganciclovir immediately after infection and cell-free supernatant collected at 3dpi. Infectious virus was quantified by plaque assay as above.

To determine if the fibrin gel matrix is inhibitory to viral release (or infection), we also infected fibroblasts cultured alone in the 3D culture system (**Figure 2B**) which showed reduce infectious virus release. These data demonstrate the reduced infection seen in the skin organ culture system is likely due, at least in part, to physical restraint of the viral particles. Analysis of viral diffusion through the fibrin gel matrix (without cells, across a horizontal gradient) also shows minimal viral diffusion (data not shown).

### HCMV replication in 3D skin models can be inhibited with standard antiviral treatment

To determine if the 3D skin model supports antiviral testing, we treated fibroblasts and the 3D skin model with Ganciclovir immediately following infection, and harvested supernatant at 3 days post-infection for measurement of infectious virus by plaque assay. As shown in **Figure 2C**, Ganciclovir inhibits viral replication in both fibroblast monolayers and 3D multicellular systems.

## Methods

### Human cytomegalovirus (HCMV)

HCMV used here is the bacterial artificial chromosome (BAC) clone of a low-passage clinical strain (TB40/E) that contains a constitutively expressed GFP for detection of viral infection (11-13). All virus was reconstituted, expanded, and titered by plaque assay on normal human dermal fibroblasts (NHDFs). Virus was used at passage 7 or lower for all studies.

### Monolayer cultures

Normal human dermal fibroblasts (NHDFs) were cultured in DMEM with 10% FBS and 1% penicillin/streptomycin/glutamine. Keratinocytes (HaCaT cells) were cultured in low calcium DMEM containing 10% FBS and 1% penicillin/streptomycin/glutamine. All cells were cultured in 5% CO_2_ at 37°C. All monolayer cultures were grown in standard tissue culture treated flasks and/or plates.

### 3D biofabricated skin models

A bioink of the fibroblast-laden dermal matrix (10^5^ cells/mL human dermal fibroblasts, 10mg/mL fibrinogen, 1 U/mL thrombin, 0.025mg/mL aprotinin, and 10% v/v glycerol in Ca^2+^-free DMEM) was added to each well of a 24 well plate. Four ∼1.5uL hemispherical fibrin droplets each seeded with 1.5 × 10^3^ HaCaT cells were printed onto the supporting matrix at approximately 2mm center-to-center spacing using a custom-built 3D bioprinter as previously described (8). Multicellular tissue constructs were cultured in aprotinin-supplemented Ca^2+^-free DMEM for ∼4-7 days to allow cell adaptation to the hydrogel matrix and keratinocyte proliferation to initiate collective migration out of the source droplets. 25µg/mL aprotinin was used to stabilize the dermal matrix that supports keratinocyte migration. 3D models and control cells were then transported between labs at ambient temperature with no more than 30 min in transit.

### HCMV infection

Cultures were infected with TB40/E-GFP pre-titered on fibroblasts by standard plaque assay for 4 hours, washed twice with PBS, and replaced with fresh complete, low-serum (2%) media. Mock-infected cultures were treated in parallel. As the number of cells in each skin model well can vary due to proliferation and differentiation in the model, infection was performed in parallel with fibroblasts and the exact infectious dose back-calculated to MOI for each experiment. Infection was performed in low serum-media on day 10-14 post fabrication using DMEM with 2% FBS, 1% penicillin/streptomycin/glutamine for 4 hours. Cells for infection experiments were cultured in DMEM with 2% FBS, 1% penicillin/streptomycin/glutamine, 25µg/mL aprotinin, and 1.6mM Ca^2+^ (introduced for the maturation of cell-cell adhesion in epidermal layers).

### Harvest of samples

Monolayer cultures were harvested by scraping and centrifugation separation of cell and supernatant fractions. 3D cultures were harvested by removal of supernatant and incubation with Nattokinase to dissolve the fibrin gel and release cells. Both supernatant fraction and dissolved gel were separated by centrifugation to achieve the standard cell and supernatant fractions comparable to monolayer cultures. Harvest timepoints are shown with each figure.

### Quantification of infectious virus

Superntant or resuspended cell fractions from growth curves were plated in duplicate on confluent monolayer cultures of NHDFs. After 2 hours, cells were washed twice with PBS and media replaced with DMEM containing 2% FBS, 1% penicillin/streptomycin/glutamine and carboxymethecellulose solution. Plaques were counted using standard methods at two weeks post-infection. Data is shown as mean with standard error of the mean for separate experimental replicates.

### Imaging

Cellular morphology and GFP positive cells were visualized with a SpectraMax MiniMax 300 Imaging Cytometery (Molecular Devices) allowing the same plate coordinates to be imaged sequentially. Transmitted light and GFP images were merged using ImageJ. GFP was used as a reporter for viral infection. SpectraMax settings were adjusted at each experiment then unmodified for all images (groups and timepoints). Images were processed only to merge GFP and transmitted light in ImageJ.

## Statistical analysis

All analyses were performed using Prism v9 as described in the figure legends.

## Discussion

This study demonstrates that a biofabricated multicellular 3D skin culture system containing both fibroblasts and keratinocytes can be infected with HCMV (**Figure 1**) and maintained for longitudinal analysis over at least 14 days (**Figure 1**). Additionally, viral replication is MOI dependent, permissive for production of infectious virus particles (**Figure 2A**), and recapitulates suppression of viral replication upon treatment with Ganciclovir (**Figure 2C**). HCMV is readily detectable in the 3D skin model and provides a tractable experimental system in which infection can be studied in a tissue-like architecture rather than exclusively in a monolayer. This observation is consistent with previous evidence that HCMV can infect multiple cell types within human skin. A prior study demonstrated that HCMV infection in human skin explants is predominantly in the dermis, including fibroblasts and endothelial cells (2) and clinical case reports of cutaneous HCMV infection demonstrate abnormal endothelial, epithelial, and fibroblast pathology (14-16). Fibroblasts, epithelial cells, endothelial cells, and smooth muscle cells, are also targets of HCMV infection in systemic organs (e.g., gastrointestinal tissue, lung) (17). Keratinocytes have a more nuanced infection profile, but more recent evidence suggests that they are involved at the oral cavity as sites of primary infection and that cellular differentiation is intrinsically linked both to infection and host responses (18, 19).

An important feature of the current system is the analysis of infection within an organized multicellular structure. Prior work (8) demonstrates that this model recapitulates stratified epidermal layers, including the basal-to-suprabasal transition. Additionally, the bioprinting model is tractable and expandable, allowing incorporation of other cell types (e.g., epithelial and endothelial cells) and control of cellular differentiation by manipulation of culture conditions.

Further improvements, including incorporation of vasculature to the model would allow study of infection spread and analysis in an advanced system, particularly by inclusion of immune cell populations (e.g., monocytes) and their contribution. Our study demonstrates that HCMV can infect and replicate in a spatially organized multicellular culture that can be experimentally manipulated and monitored longitudinally. Additionally, both viable cells (data not shown) and infectious virus can be recovered from the biofabrication matrix (data not shown) and infectious virus recovered from the extracellular space (**Figure 2**) demonstrating feasibility for additional studies. However, several limitations remain. We have not yet determined the identity of infected cells nor their spatial relationship to each other within the culture beyond what is shown (**Figure 1** and **Supplemental Figures 1-3**).

These points highlight the value of this system for future studies, and our data demonstrates that the system is tractable, expandable, and supports productive viral replication, demonstrating that it can be used to test the contribution of cell populations to HCMV spread, whether tissue architecture changes the balance between cell-associated and extracellular virus, and whether host inflammatory or differentiation states alter viral replication in a multicellular system. The current biofabricated model is less physiological complex but offers greater control over cellular composition and experimental conditions than human skin organ cultures. In parallel, which conventional 2D cultures provide greater simplicity and quantitative analysis, they lack the spatial organization of the biofabricated model or skin explants. In summary, this system may provide a useful intermediate between monolayer cultures and ex vivo or in vivo human tissues, as this skin model supports productive viral infection, including release of infectious virus and suppression of viral replication via Ganciclovir treatment, and provides a tractable and spatially organized multicellular 3D environment for future studies on viral replication, spread, host responses, and tissue architecture.

## Supporting information

all supplemental figure panels

## Author Contributions

HZ: investigation, writing – review and editing. HN, OW, MB: investigation PA: formal analysis, writing – review and editing, funding acquisition. FM: formal analysis, writing – review and editing, funding acquisition, supervision. LBC: investigation, formal analysis, writing – original draft, writing – review and editing, funding acquisition, supervision.

---

**Supplemental Figure 1. Longitudinal analysis of replicate fibroblast monolayer cultures**. Monolayer cultures of fibroblasts (as in Figure 1) were infected with HCMV and imaged using a SpectraMax Imaging Cytometer (Molecular Devices) allowing remeasurement of the same coordinates on day 0, 3, 5, 7, 10, and 14dpi. Both infected (HCMV) and uninfected (Mock) controls are shown for both transmitted light and GFP expressing cells. Experiment done in parallel with Supplemental Figures 2 and 3.

**Supplemental Figure 2. Longitudinal analysis of replicate keratinocyte monolayer cultures**. Monolayer cultures of keratinocytes (as in Figure 1) were infected with HCMV and imaged using a SpectraMax Imaging Cytometer (Molecular Devices) allowing remeasurement of the same coordinates on day 0, 3, 5, 7, 10, and 14dpi. Both infected (HCMV) and uninfected (Mock) controls are shown for both transmitted light and GFP expressing cells. Experiment done in parallel with Supplemental Figures 1 and 3.

**Supplemental Figure 3. Longitudinal analysis of replicate 3D skin organ multi-cellular cultures**. 3D cultures of fibroblasts and keratinocytes (as in Figure 1) were infected with HCMV and imaged using a SpectraMax Imaging Cytometer (Molecular Devices) allowing remeasurement of the same coordinates on day 0, 3, 5, 7, 10, and 14dpi. Both infected (HCMV) and uninfected (Mock) controls are shown for both transmitted light and GFP expressing cells. Experiment done in parallel with Supplemental Figures 1 and 2.

## References

1. Howley PM, Knipe DM, Cohen JL, Damania BA. 2021. Fields Virology: DNA Viruses, vol 2. Wolters Kluwer.

2. Lloyd MG, Smith NA, Tighe M, Travis KL, Liu D, Upadhyaya PK, Kinchington PR, Chan GC, Moffat JF. 2020. A Novel Human Skin Tissue Model To Study Varicella-Zoster Virus and Human Cytomegalovirus. J Virol 94:JVI.01082-20.

3. Morrison KM, Beucler MJ, Campbell EO, White MA, Boody RE, Wilson KC, Miller WE. 2019. Development of a Primary Human Cell Model for the Study of Human Cytomegalovirus Replication and Spread within Salivary Epithelium. J Virol 93.

4. Brown RM, Rana P, Jaeger HK, O’Dowd JM, Balemba OB, Fortunato EA. 2019. Human Cytomegalovirus Compromises Development of Cerebral Organoids. J Virol 93.

5. Egilmezer E, Hamilton ST, Foster CSP, Marschall M, Rawlinson WD. 2024. Human cytomegalovirus (CMV) dysregulates neurodevelopmental pathways in cerebral organoids. Communications Biology 7:340.

6. Sun G, Chiuppesi F, Chen X, Wang C, Tian E, Nguyen J, Kha M, Trinh D, Zhang H, Marchetto MC, Song H, Ming G-L, Gage FH, Diamond DJ, Wussow F, Shi Y. 2020. Modeling Human Cytomegalovirus-Induced Microcephaly in Human iPSC-Derived Brain Organoids. Cell Reports Medicine 1.

7. Ijezie EC, O’Dowd JM, Kuan MI, Faeth AR, Fortunato EA. 2023. HCMV Infection Reduces Nidogen-1 Expression, Contributing to Impaired Neural Rosette Development in Brain Organoids. Journal of Virology 97:e01718–22.

8. Zhai H, Jin X, Minnick G, Rosenbohm J, Hafiz MAH, Yang R, Meng F. 2022. Spatially Guided Construction of Multilayered Epidermal Models Recapturing Structural Hierarchy and Cell–Cell Junctions. Small Science 2:2200051.

9. Zhai H, Jin X, Biegert M, Zahs B, Yang R, Meng F. 2026. Stepwise Generation of Vascularized Multilayered 3D Organotypic Skin Models. Bio-protocol 16:e5802.

10. Zhai H, Jin X, Moghaddam AO, Spieker MF, Seiffert-Sinha K, Sinha AA, Yang R, Meng F. 3D organotypic skin models recapitulate autoantibody-driven pemphigus pathomechanisms and targeted therapeutic response. Science Advances 12:eadz5003.

11. Crawford LB, Caposio P, Kreklywich C, Pham AH, Hancock MH, Jones TA, Smith PP, Yurochko AD, Nelson JA, Streblow DN. 2019. Human Cytomegalovirus US28 Ligand Binding Activity Is Required for Latency in CD34(+) Hematopoietic Progenitor Cells and Humanized NSG Mice. mBio 10.

12. Hancock MH, Crawford LB, Perez W, Struthers HM, Mitchell J, Caposio P. 2021. Human Cytomegalovirus UL7, miR-US5-1, and miR-UL112-3p Inactivation of FOXO3a Protects CD34(+) Hematopoietic Progenitor Cells from Apoptosis. mSphere 6.

13. Crawford LB. 2022. Human Embryonic Stem Cells as a Model for Hematopoietic Stem Cell Differentiation and Viral Infection. Curr Protoc 2:e622.

14. Choi YL, Kim JA, Jang KT, Kim DS, Kim WS, Lee JH, Yang JM, Lee ES, Lee DY. 2006. Characteristics of cutaneous cytomegalovirus infection in non-acquired immune deficiency syndrome, immunocompromised patients. British Journal of Dermatology 155:977–982.

15. Drozd B, Andriescu E, Suárez A, De la Garza Bravo MM. 2019. Cutaneous cytomegalovirus manifestations, diagnosis, and treatment: a review. Dermatol Online J 25.

16. Odani K, Itoh A, Yanagita S, Kaneko Y, Tachibana M, Hashimoto T, Tsutsumi Y. 2020. Paraneoplastic Pemphigus Involving the Respiratory and Gastrointestinal Mucosae. Case Rep Pathol 2020:7350759.

17. Sinzger C, Grefte A, Plachter B, Gouw AS, The TH, Jahn G. 1995. Fibroblasts, epithelial cells, endothelial cells and smooth muscle cells are major targets of human cytomegalovirus infection in lung and gastrointestinal tissues. J Gen Virol 76 (Pt 4):741–50.

18. Cojohari O, Cao Y, Temirbek A, Garber M, Colubri A, Kowalik T. 2025. Human Cytomegalovirus Infection of Primary Human Oral Keratinocytes Induces Intermediate Keratinocyte Differentiation and an Altered Innate Immune Response. bioRxiv doi:10.1101/2025.09.16.676604.

19. Weng C, Lee D, Gelbmann CB, Van Sciver N, Nawandar DM, Kenney SC, Kalejta RF. 2018. Human Cytomegalovirus Productively Replicates In Vitro in Undifferentiated Oral Epithelial Cells. J Virol 92

