## Supplementary figures and images for "Human Cytomegalovirus Infects and Remodels 3D-biofabricated Skin Equivalents"

### all supplemental figure panels

*Supplemental Figure 1*

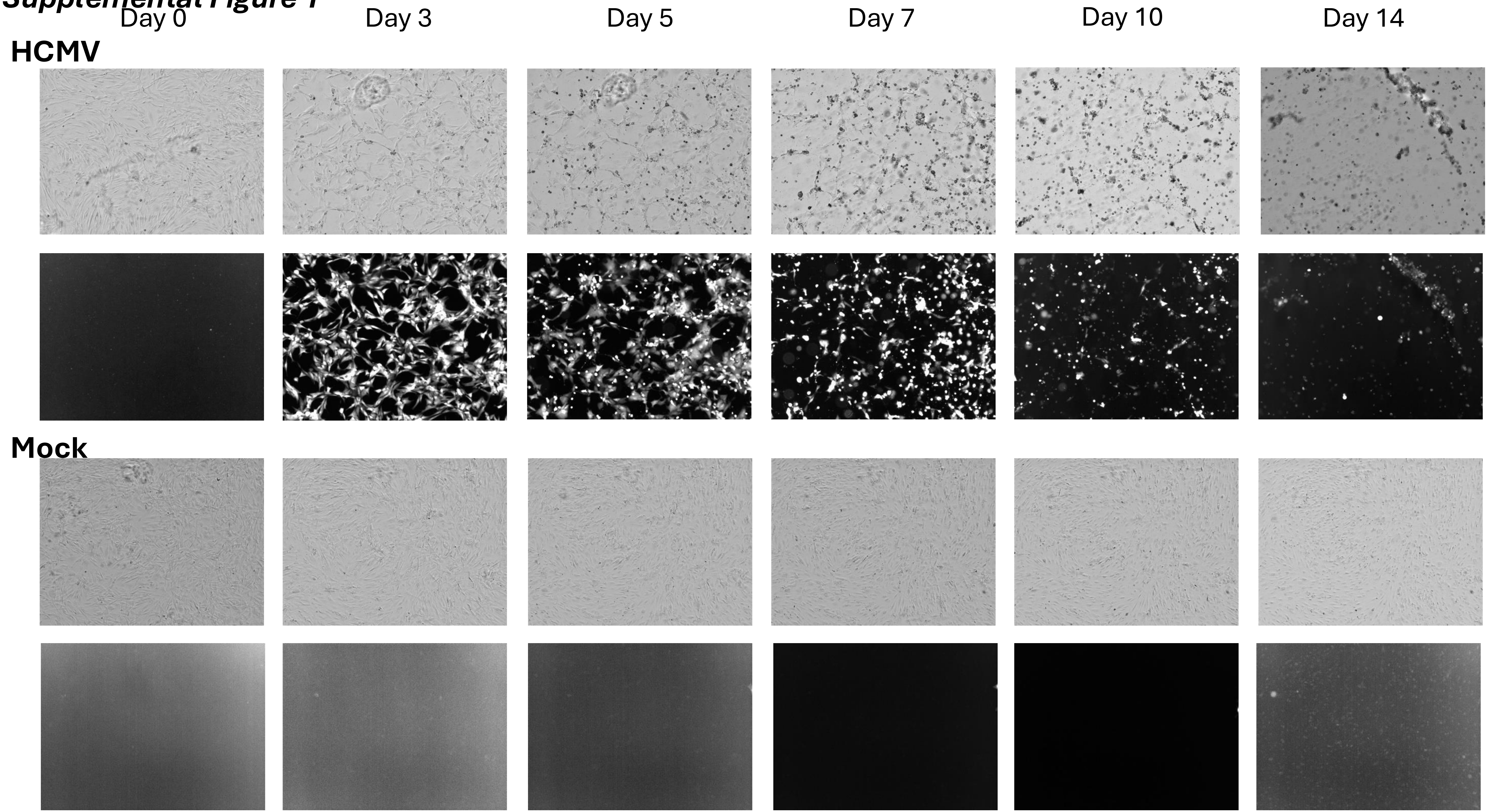

*Supplemental Figure 2*

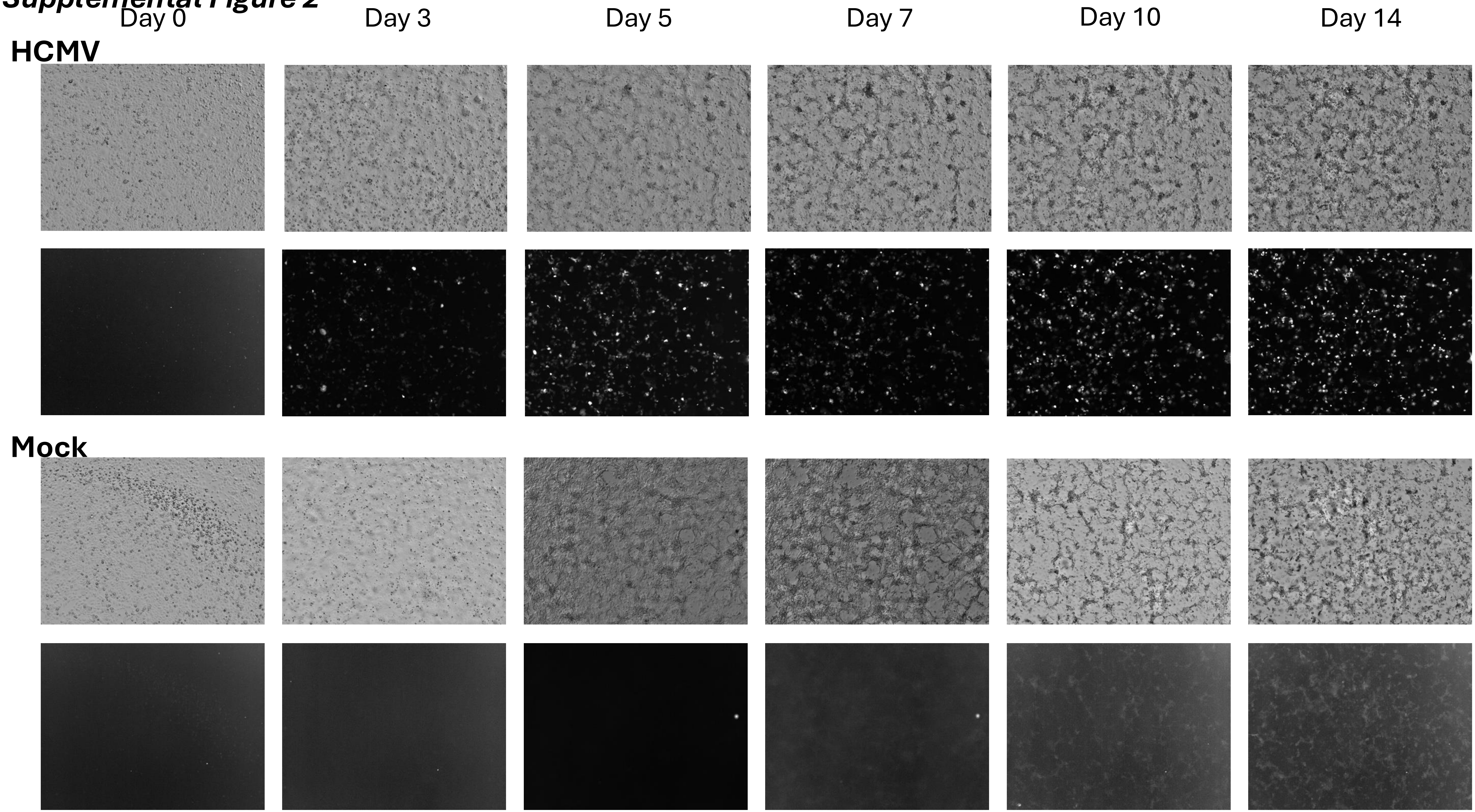

***Supplemental Figure 3***

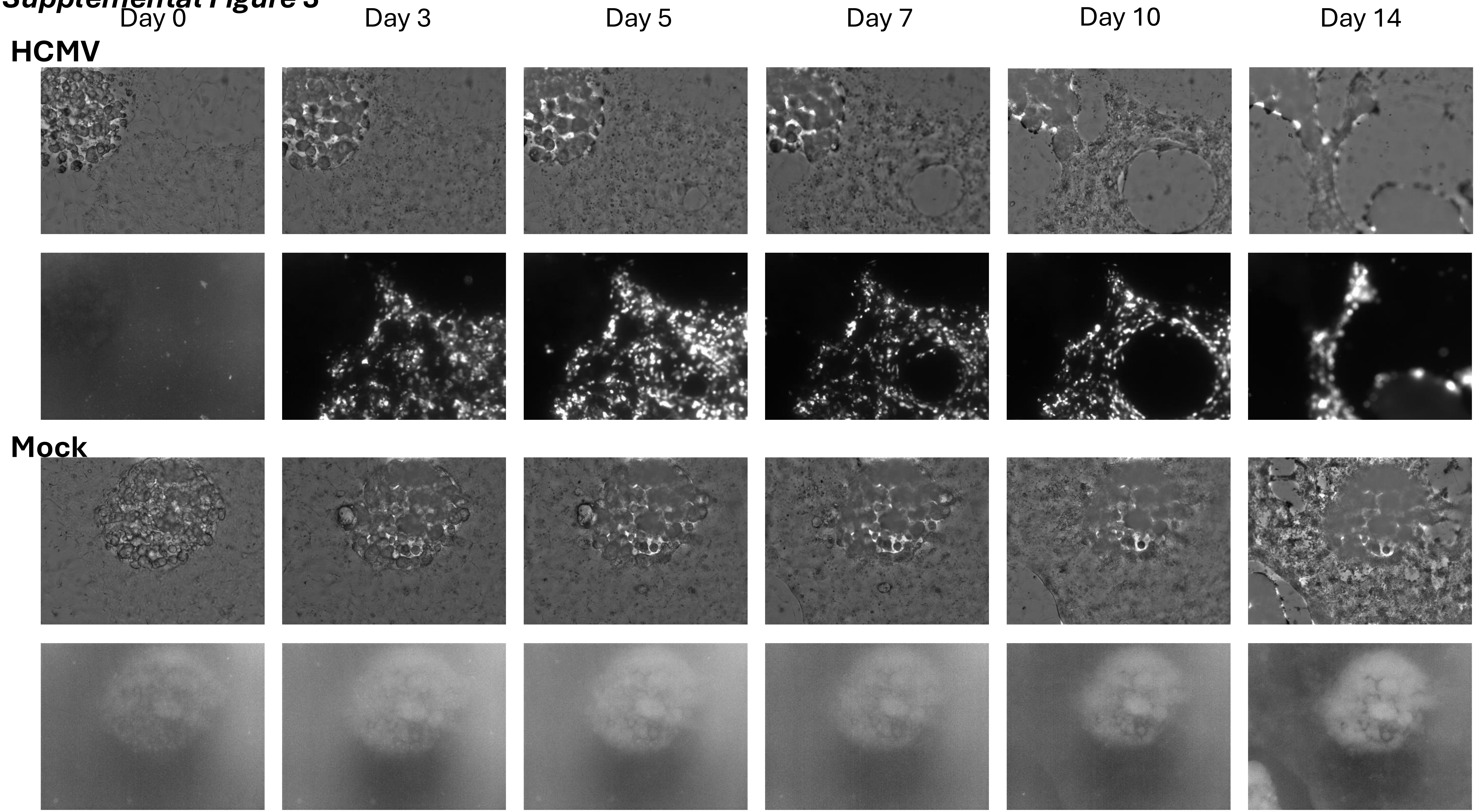
